# The Blueprint for Survival: The Blue Dasher Dragonfly as a Model for Urban Adaptation

**DOI:** 10.1101/2024.12.12.628234

**Authors:** Ethan R. Tolman, Ellie Gamett, Christopher D. Beatty, Aaron Goodman, Brittney Hahn, Christian Benischek, Gracie Castillo, Ethan Derderian, Santiago Fernandez-Juarez, Ben Gallafent, James Jenson, Dick Jordan, Magnolia Schneider, Roberto Salazar, Towako Tamano, Maleah Wei, Jacob Idec, Rob Guralnick, Jessica L. Ware, Manpreet K. Kohli

## Abstract

Human alteration of natural environments and habitats is a major driver of species decline. However, a handful of species thrive in human altered environments. The biology, distribution, population structure, and molecular adaptations enabling certain species to thrive in human-altered habitats are not well understood. Here, we evaluate the population and functional genomics, ecological niche and distributions, and geometric morphometrics of the blue dasher (*Pachydiplax longipennis*), one of the most ubiquitously observed insects in human altered habitats. Using resequencing data we identify a number of genes involved with the success of the blue dasher in human altered habitats, including loci contributing to immune function and response to oxidative stress. Some genes related to these functions are found in regions of strong population structure, while others are not, potentially indicating both regional and widespread adaptations to urban environments within this species. Using one of the most robust locality datasets for any species to date, we also generate habitat suitability predictions which show that *P. longipennis* has spread with urbanization, suggesting humans have created suitable habitat for this species. These results complement morphological and genomic data showing *P. longipennis* (particularly East of the Rocky Mountains) has the capacity to rapidly disperse to newly suitable habitats. Given the shared barriers to colonizing an urban habitat, we expect that many of the adaptations we have identified in *P. longipennis* can be used for predicting what animals might succeed in urban habitats more generally.

## Introduction

In the Anthropocene, population decline is the norm for many species [1,2]. However, some species are thriving on our changing planet. One such “winner” of the Anthropocene is the blue dasher dragonfly, *Pachydiplax longipennis* (Brauer 1868). Not only is the range of this species thought to be expanding with a warming climate [3], gene family expansions related to the processing of free radicals and immune function are hypothesized to allow this species to live in highly polluted lentic bodies of water near humans [4]. As a highly photogenic “percher” species which spends a great deal of its time perched on vegetation [5,6], *P. longipennis* is easily detectable through community science tools, such as iNaturalist. Due to its proximity to urban habitats, *P. longipennis* was the most observed dragonfly on iNaturalist through December 2024 and the 40th most observed species in its distribution. As a robust generalist with the potential to displace specialists [4,7], a comprehensive analysis of this species is important for the monitoring of aquatic insect communities on a rapidly changing planet, yet no work has been done to explore regional genetic, genomic and phenotypic differences in *P. longipennis*. Here we consider the life history, population and functional genomics, geometric morphometrics, and ecological niche of *P. longipennis* populations in North America to test the hypothesis that this species is expanding its range and is equipped for colonizing newly altered habitats where more specialist species are unable to persist. We also identify loci involved in success in urban habitats, and more generally, make a case for establishing *P. longipennis* as an indicator of altered habitats.

### Pachydiplax longipennis life history

*P. longipennis* is distributed across North America, from southern Canada in the north, to Mexico and Belize in the south, and is found on either side of the Rocky Mountains [6,8]. It is also found on the Caribbean islands of Cuba and the Bahamas, as well as Bermuda, although it is currently unclear when or how it arrived on these islands [8]. Blue dashers are medium to small dragonflies in the family Libellulidae, roughly 2.5 to 4.5 cm in length, with striped thoraces, dark colouration at the base of their wings, and waxy blue pruinosity in mature males [6]. *Pachydiplax* is a monotypic genus that has been consistently recovered in a clade that includes the genera *Micrathyria* and *Zenithoptera* [9–11]. The features of males, including their size and dispersal [12,13], aggressiveness, feeding and territoriality [14–18], and nymph microhabitat preference and emergence details [19], have been well studied, but data on females is generally lacking.

*P. longipennis* has long been considered a potential migrant species, as it has been found in swarms of migrating *Anax junius* on several occasions [20–22], and was considered a putative migrant by the (no longer active) Migratory Dragonfly Partnership [23]. Recently, individuals of *Pachydiplax longipennis* were observed in a large migratory mixed swarm of dragonflies that flew over Rhode Island in July of 2024 [20]. Migratory, or even high dispersing, behavior may allow *P. longipennis* to more rapidly colonize newly suitable habitat, which has been observed anecdotally [20,22], but no molecular or morphological study has been used to assess gene flow within, or the migratory capabilities of, this species.

## Materials and Methods

### Population genomics

We obtained low coverage 150 bp Illumina data from Azenta Life Sciences for 15 individuals, five each from Boise, Idaho; Newark, New Jersey; and Knoxville, Tennessee. We mapped the trimmed reads, delivered from the manufacturer, with BWA [24], to the previously assembled reference genome of *P. longipennis* [4], using default settings. We marked and removed duplicates with Picard [25], and called variants with bcftools v1.15.1 [26], using the options: `bcftools mpileup-a AD,DP,SP-Ou- f reference.fasta. Input_bam | bcftools call-f GQ,GP-mO z-o output`. We filtered the resulting vcf file for a minimum quality score of 30, and a minimum and maximum depth of 1 and 6 respectively. We utilized PLINK v1.90b6.21 [27] to prune linked sites. To estimate population structure, we performed a principal components analysis (PCA) and calculated the F_ST_ between and within each of the three populations using the linkage-pruned SNPs with VCFtools v0.1.16 [27]. To estimate the ancestry in each of our populations, we generated admixture models assuming 2-5 ancestral populations with ADMIXTURE v1.3.0 [28] and evaluated the models with cross-validation error. The best fitting admixture-model and the PCA plots were plotted in R using mapmixture v1.1.0 [29].

In addition to the called SNPs, we also estimated population structure analyses from genotype likelihoods. We used ANGSD v.0.921 [30] to calculate genotype likelihoods with the options “- remove_bads 1-uniqueOnly 1-only_proper_pairs 1-C 50-baq 1-minMapQ 30-minQ 30-minInd 13-doSnpStat 1-doHWE 1-sb_pval 1e-6-hwe_pval 1e-6-hetbias_pval 1e-6-doMajorMinor 1-skipTriallelic 1-doMaf 2- doPost 1-minMaf 0.05-snp_pval 1e-6-doGeno 8-doGlf 2”. We then used ngsLD v1.2.1 [31] to identify unlinked sites, which were used as input for PCangsd [32], to evaluate population structure through a PCA. We also estimated admixture models from the genotype likelihoods using NGSadmix [28], with 2-5 ancestral populations. As with the SNP analyses, both the principal components and admixture analyses were plotted in R using mapmixture [29].

To identify specific genomic regions and genes of high structure between our three populations, we used VCFtools [26] to calculate Fst across a sliding window of 10,000 bp, with a step size of 1,000bp. We then identified genes, (previously annotated by Tolman et al. [33]) in windows of high (F_ST_ >.5) and low (F_ST_ <.01) structure between populations and used REVIGO [34] to find and summarize the molecular functions of the genes in these regions into treemaps, using the previously published functional annotations [4].

### Demographic modeling

To model historical gene flow between our sampled populations of *P. longipennis* we generated a demographic model for *P. longipennis* using GADMA2 v2.0.0 [35]. We used the LD-trimmed SNP file as input with the default engine moments [36], a mutation rate of 2e-9 (previously shown to be a reasonable estimate for Odonata [37]), and the local optimizer BFGS_log (see supplementary materials for full configuration file).

### COI Haplotype Structure

To broaden our population sampling we downloaded all COI sequences for *P. longipennis* from BOLD [38], and also extracted the COI sequence from the reference assembly of *P. longipennis* [33], which was generated from one individual from Boise, Idaho. We also obtained Sanger sequencing (performed by Genewiz) for the COI gene (using primers and protocol as specified in Kohli et al [39]) for samples collected in New Jersey and Tennessee. The sequences were aligned in CLUSTAL OMEGA (online server https://www.ebi.ac.uk/jdispatcher/msa/clustalo) [40] and then manually corrected in MESQUITE v1.8 [41]. All together we had sequences from the following locations: Florida (24), Ontario (7), Arkansas (9), Texas (5), Arizona (11), Oklahoma (1), British Columbia (1), New Jersey (19), Idaho (1), and Tennessee (2). The population structure and the corresponding haplotype network and population statistics for COI were estimated in software package PopART [42]. To test for population structure on either side of the Rocky Mountain divide, we considered Arizona, Idaho, and British Columbia to belong to a “West Coast” population, with the remaining individuals belonging to an “East Coast” population, and used PopART [42] to estimate the COI F_ST_ between the West and East coast populations. As the total sequence length for two Tennessee individuals was less than other samples, we performed all our COI analyses in PopART both with and without the individuals from Tennessee.

### Morphological analysis

To investigate regional morphological differences in the wings of *P. longipennis*, which can have a major effect on dragonfly dispersal [43], we collected specimens from two locations: Boise, Idaho and Herndon, Virginia and compared them to museum specimens from Secaucus, New Jersey and Knoxville, Tennessee. We collected a total of 28 individuals from the combined four locations (14 from Idaho, 6 from Virginia, 9 from Tennessee, and 10 from New Jersey). Live specimens from Virginia and Idaho were carefully secured by their wings to labeled index cards using paper clips to avoid any damage to the dragonflies. Each individual was photographed alongside a ruler for scale reference (Fig. 1). Museum specimens were scanned with an Epson Perfection V550 Photo flatbed color scanner according to a digitization template created for the Targeted Odonata Wing Digitization Project v2.0 (Supplementary figure 1). Specimens were imaged with the ventral side of the right pair of wings facing down on the glass.

**Figure 1:**
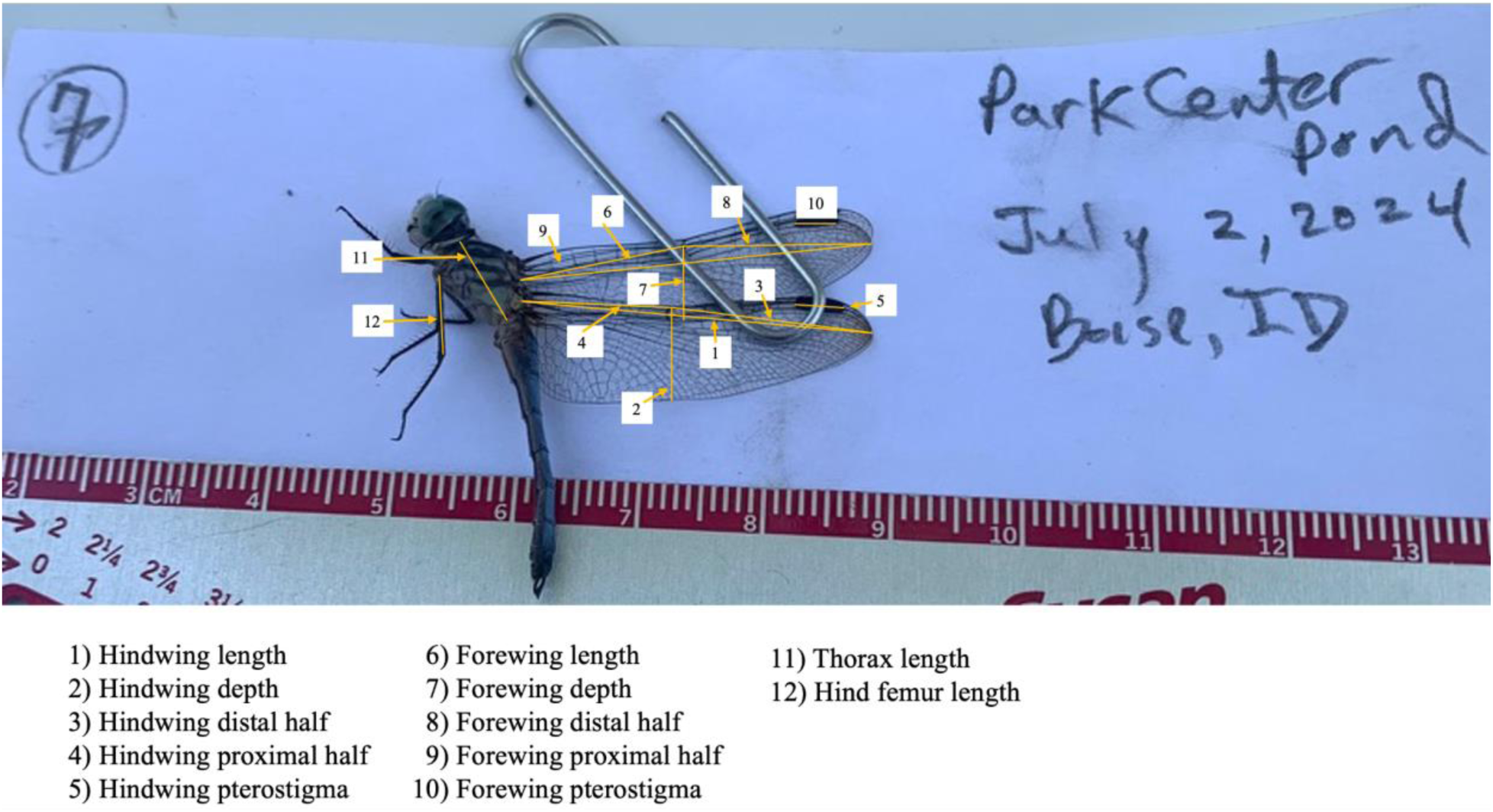
Morphological measurements taken of P. longipennis. An example of a photograph taken for measurements, with the measurements labeled on the figure.

Four researchers then independently measured the hindwing length, hindwing depth, hindwing pterostigma length, length of the hindwing distal half (wing node to tip), length of the hindwing proximal half (wing node to base), forewing length, forewing depth, forewing pterostigma length, length of the forewing distal half, length of the forewing proximal to the body, thorax length and femur length (Fig. 1) in ImageJ [44]. An average of each measurement taken by the four researchers was used in statistical analyses. We attempted to capture the area of the fore and hindwings using the protocol established by Idec et al. [45].

To determine whether there were significant morphological differences between the four geographic populations (Idaho, Virginia, Tennessee, and New Jersey) that may affect the dispersal capability of *P. longipennis*, we applied the Kruskal-Wallis test in R v4.1.4 [46] by scaling each wing measurement according to thorax length and femur length to control for differing wing measurements being a result of a larger body size in general between the populations [47,48]. This non-parametric method is suitable for comparing more than two groups, as it does not assume a normal distribution. Variables with p-values below 0.05 were considered to show significant regional differences. As our gender sampling was uneven (Idaho were all males, all but one individual from New Jersey were females, and the other populations were mixed) we compared males and females in a combined sample of the three genetically similar east coast populations with a Wilcoxon rank-sum test. As we determined that our wing measurements differed by sex when normalized by thorax length, but not when normalized by femur length, we considered measurements normalized by femur length in data visualization and further analyses. To further visualize the morphological space inhabited by each population we conducted a PCA. We imputed 26 missing measurements for each population using the package mice v3.16.0 [49] and conducted a PCA using the wing measurements normalized by femur length in R [46]. We visualized the PCA with mapmixture v1.1.4 [29]

### Ecological niche modeling

#### Occurrence Data

We selected occurrences for *P. longipennis* using the Global Biodiversity Information Facility (GBIF), pulling data from citizen science sources and natural history museum collections (iNaturalist, Odonata Central, National Museum of Natural History). We selected research-grade occurrences with verified latitude and longitude coordinates, a photograph of the sighting, observation date, and at least ⅔ agreement on species identification by the community. Furthermore, we filtered occurrences by removing erroneous localities (natural history museums, middle of the ocean, etc.). We binned our occurrences to 5- year intervals between the years 1990-2020, which consisted of 6 time intervals, each with differing numbers of occurrences:

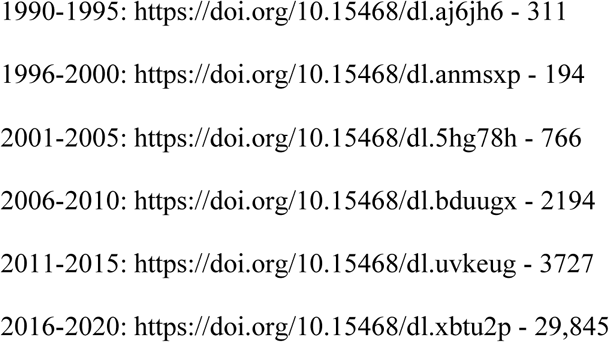

#### Environmental Data

All modeling and environmental raster processing was conducted using the statistical program R v. 4.1.2 with the packages ENMeval v2.0 [50,51], WALLACE v.1.9 [52], and dismo [53]. We acquired monthly minimum and maximum temperature and precipitation rasters of the United States from 2000-2020 using the Worldclim “Historical Monthly Weather Dataset” at 2.5 arcminute resolution. We averaged the monthly rasters for the 20-year interval and extrapolated the 19 bioclimatic variables using the final averages of maximum and minimum temperature and precipitation [54].

#### Model Building

We thinned our occurrence dataset to 10 km to match the resolution of our raster dataset, with the aim of filtering to 1-2 occurrences per city, and to reduce spatial autocorrelation, artificial clustering, and to even sample sizes across our time intervals. We used the spThin package [55]. We defined a study extent consisting of the contiguous United States. Within our extent, we randomly sampled 50,000 background points for modeling and extracted their environmental values. We used these values to calculate correlations between variables using the ‘vifcor’ and ‘vifstep’ functions in the ‘usdm’ package [56] and filtered out variables with correlation coefficients higher than 0.7 and a Variance Inflation Factor (VIF) threshold of 10.

MaxEnt (Maximum entropy algorithm) was chosen not only for its ability to incorporate spatial biases into the modelization process but is recommended for presence-only occurrence datasets [57–60]. MaxEnt compares the environmental conditions at locations of occurrence records with randomly selected points from a background extent to create a machine-learning model of habitat suitability. Species occurrences were spatially partitioned using the ‘checkerboard2’ method [51], whereby occurrence data is geographically split into equal partitions by overlaying two sets of checkerboards, using 25% of the localities to calibrate the models (‘training’ data) and 75% to evaluate the models’ predictive accuracy (‘evaluation’ data).

Feature classes were incorporated into the model, which determine the shape of available modeled relationships in environmental space [59]. To objectively compare modeled suitability of *P. longipennis*, and due to the high number of occurrence records (making computation with multiple feature classes unfeasible) we restricted our model relationship to linear feature classes and a regularization of 1. Both feature classes and regularization multipliers can strongly influence MaxEnt model outputs [61,62].

Optimal models were assessed using threshold independent (AUC curve) and threshold dependent metrics (10% omission). AUC (Area Under the Curve) is the measure of the ability of the MaxEnt model generated to discriminate between false positives and negatives. AUC ranges from 0 to 1, with 1 being considered a “perfect model”, 0.5 as random, and < 0.5 considered worse than random [63,64]. Optimal models were based on the averaged validation AUC, which is the mean of the AUC of the testing data for all partitions (AUC_test_). Omission rate is another method of evaluation whereby a threshold is applied to a continuous model prediction. Application of the threshold makes the model binary, in which points are categorized as either within or outside the prediction. We checked model performance using the Akaike criterion for small sample sizes (AICc) [62], as well as the number of nonzero coefficients (parameters) from each model.

For optimal models within each period, we documented the importance of variables and plotted response curves. Maxent calculates permutation importance by changing the values of each environmental variable at random, then calculating the difference using the AUC from the ‘training data’. The values of each environmental variable are randomly permuted within the training presence and background data, the resultant drop in training AUC is calculated, then normalized to percentages. The larger the percentage, the more importance that variable possesses in suitability. Response curves show how each environmental variable individually affects the maxent prediction in terms of increasing or decreasing suitability. Behavior of response curve is dictated by model complexity (feature classes and regularization multipliers) [58].

Maxent raw predictions were transformed to a scale of 0–1 to approximate probability of occurrence using the ‘cloglog’ transformation [58] and displayed on a QGIS *v.3.34*.

## RESULTS

### Sequencing and depth

With the exception of one individual from Tennessee with an average depth of 0.45, which was excluded from downstream analyses, all sequenced samples had an average sequencing depth between 2.00 and 3.40.

### Population structure

The best-fitting admixture analyses identified two ancestral populations from genotype likelihoods, and four ancestral populations from SNPs (Fig 2A-B). In the SNPs admixture analysis, ancestral populations one and three were unique to Tennessee and New Jersey (Fig. 2B). All individuals from Idaho had ancestry from ancestral populations two and four, which were also found at low levels in Tennessee (Fig. 2B). In the likelihood admixture analysis all individuals from Idaho had 100% of their ancestry assigned ancestral population two, one individual from Tennessee split its ancestry between ancestral populations one and two, and the remainder of the individuals had 100% of their ancestry assigned to ancestral population one (Fig. 2A).

**Figure 2:**
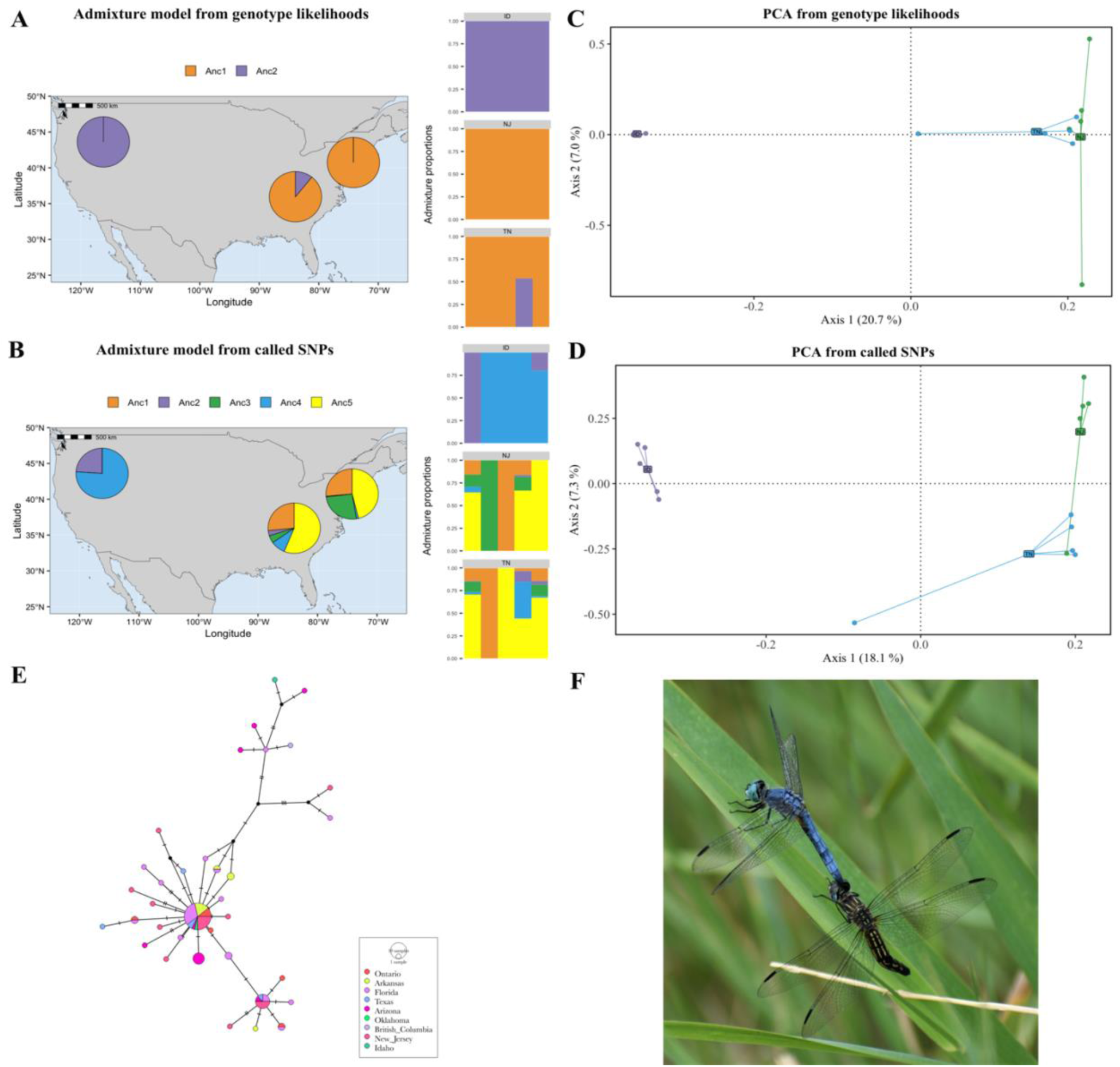
Population Structure of P. longipennis. Summary of population structure analyses calculated with whole genome resequencing data. Admixture analyses calculated using (A) genotype likelihoods, and (B) called SNPs. The best model was chosen by cross validation and is shown here. PCA plots of sampled P. longipennis calculated with (C) genotype likelihoods, and (D) called SNPs. (E) COI haplotype network of P. longipennis from publicly available COI sequences, including the COI sequence from the reference genome. (F) A male P. longipennis from Boise, Idaho mate guarding a female (credit: Kim Chmura).

The PCA, admixture, and weighted and unweighted Fst analyses, using both the called SNPs and genotype likelihoods, showed little divergence between New Jersey and Tennessee (F_ST_ <.01, Fig. 2A-D, Table 1). Between these two populations, ∼88% of genes were in regions of high genomic conservation (F_st_ < 0.01), and < 0.01% of genes were in highly structured regions (F_st_ > 0.5). None of the 31 structured genes (F_ST_ >.50) between New Jersey and Tennessee had a functional annotation, and the large number of unstructured genes between these populations was not tractable for visualization with REVIGO. The Idaho population showed comparatively average high divergence with both East Coast populations (F_ST_ > 0.18, Fig. 2A-D, Table 1). The COI haplotype analyses also showed an East/West divide, with largely shared haplotypes between East Coast populations, and differing, but closely related haplotypes in the West Coast populations (Fig. 2E), and a high F_ST_ (> 0.30) compared to the genome wide averages (Table 1). In comparing Idaho and Tennessee, we found that ∼ 31% of genes were in regions of high genomic conservation between the two populations (F_st_ < 0.01), and ∼ 9% of genes were in regions of high divergence (F_st_ > 0.5). 236 biological processes were summarized from the functional annotation of genes in highly unstructured regions between Idaho and Tennessee, including 86 cellular components and 307 molecular functions (Fig. 3). Genes from highly diverged regions of the genome were summarized into 76 biological processes, 31 cellular components, and 100 molecular functions (Fig. 3).

**Table one:**
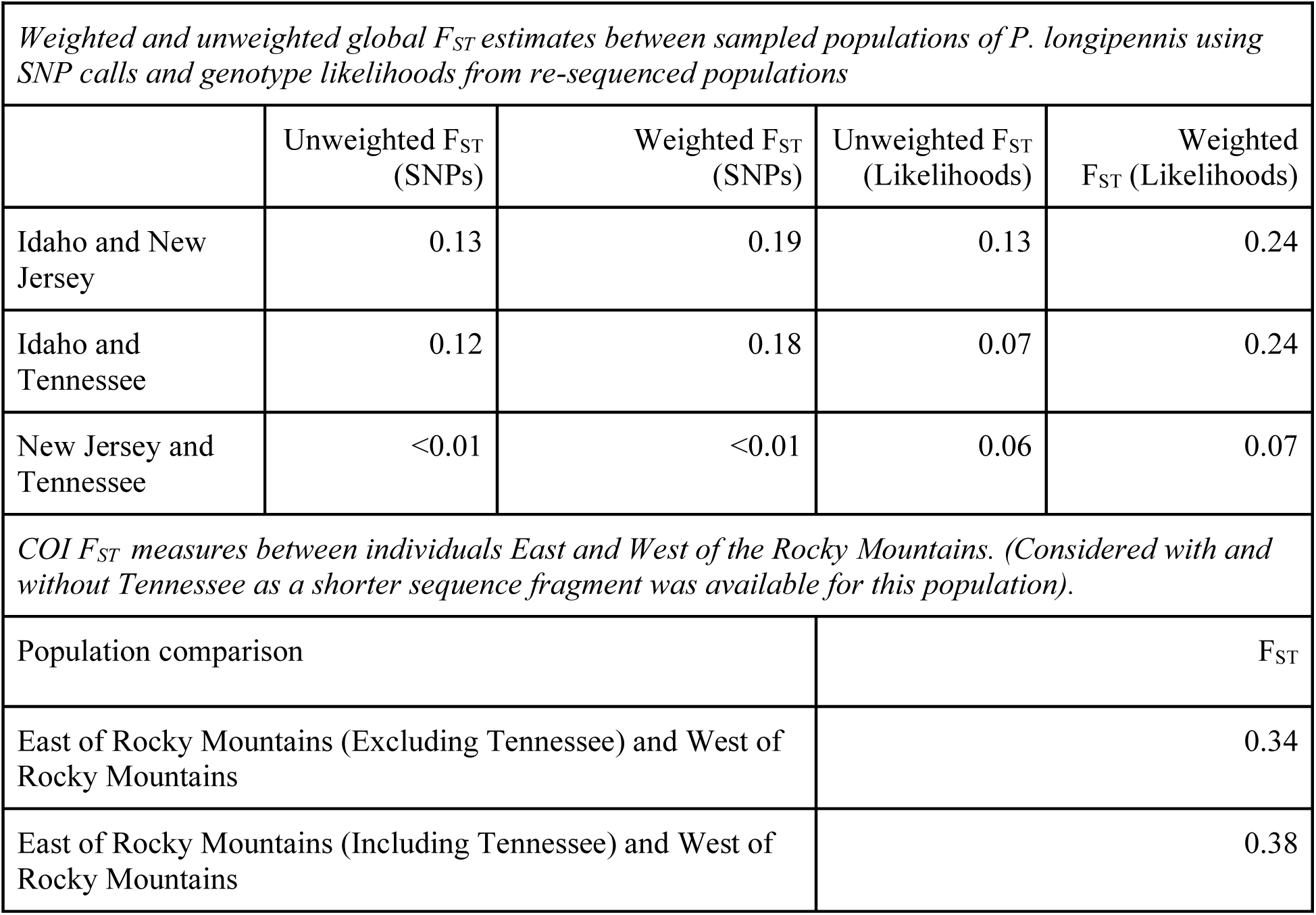
F_ST_ Estimates.

**Table three:**
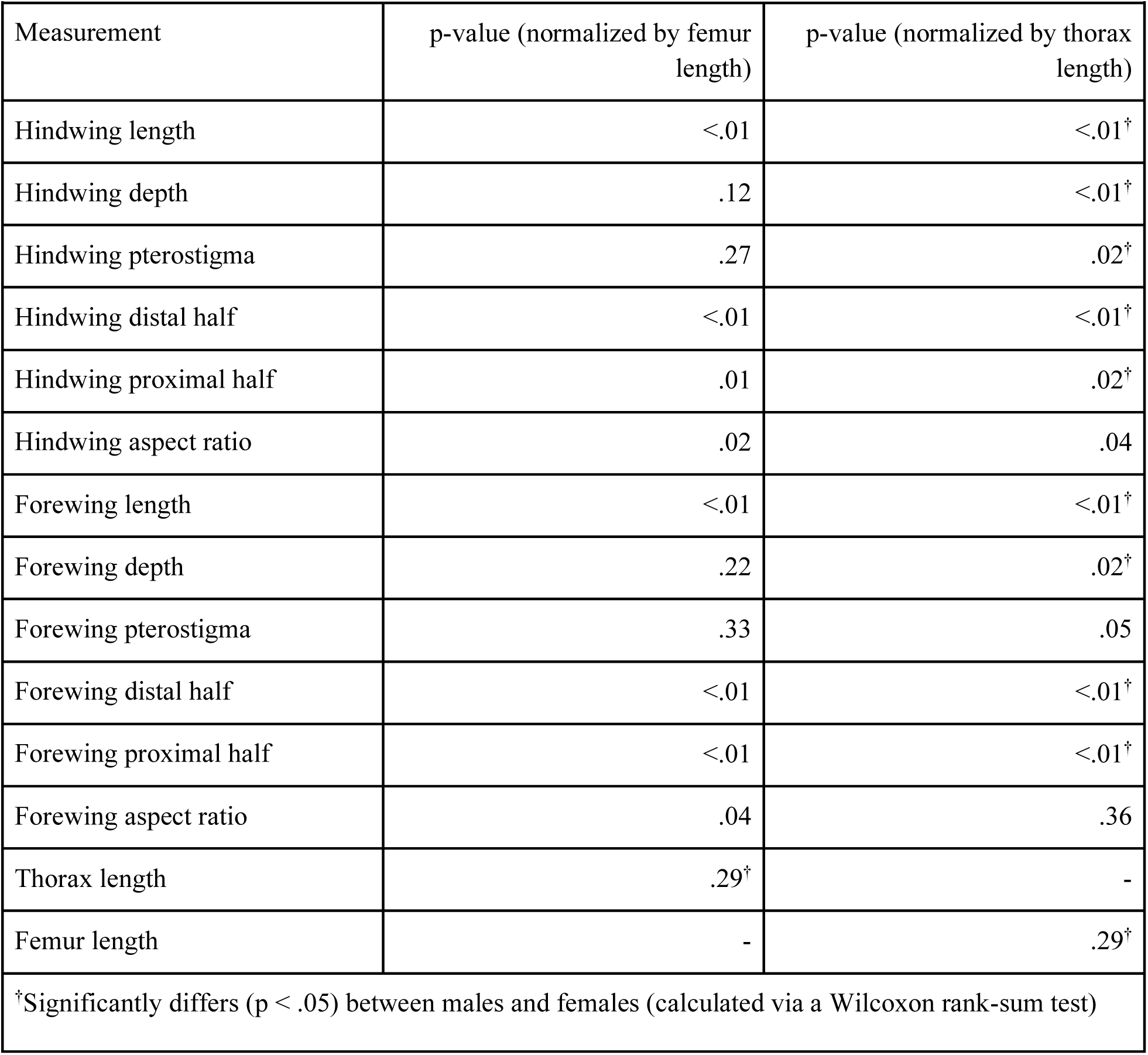
Significance of Kruskal Wallace Test for each measurement when normalized by either femur or thorax length.

**Table Four:**
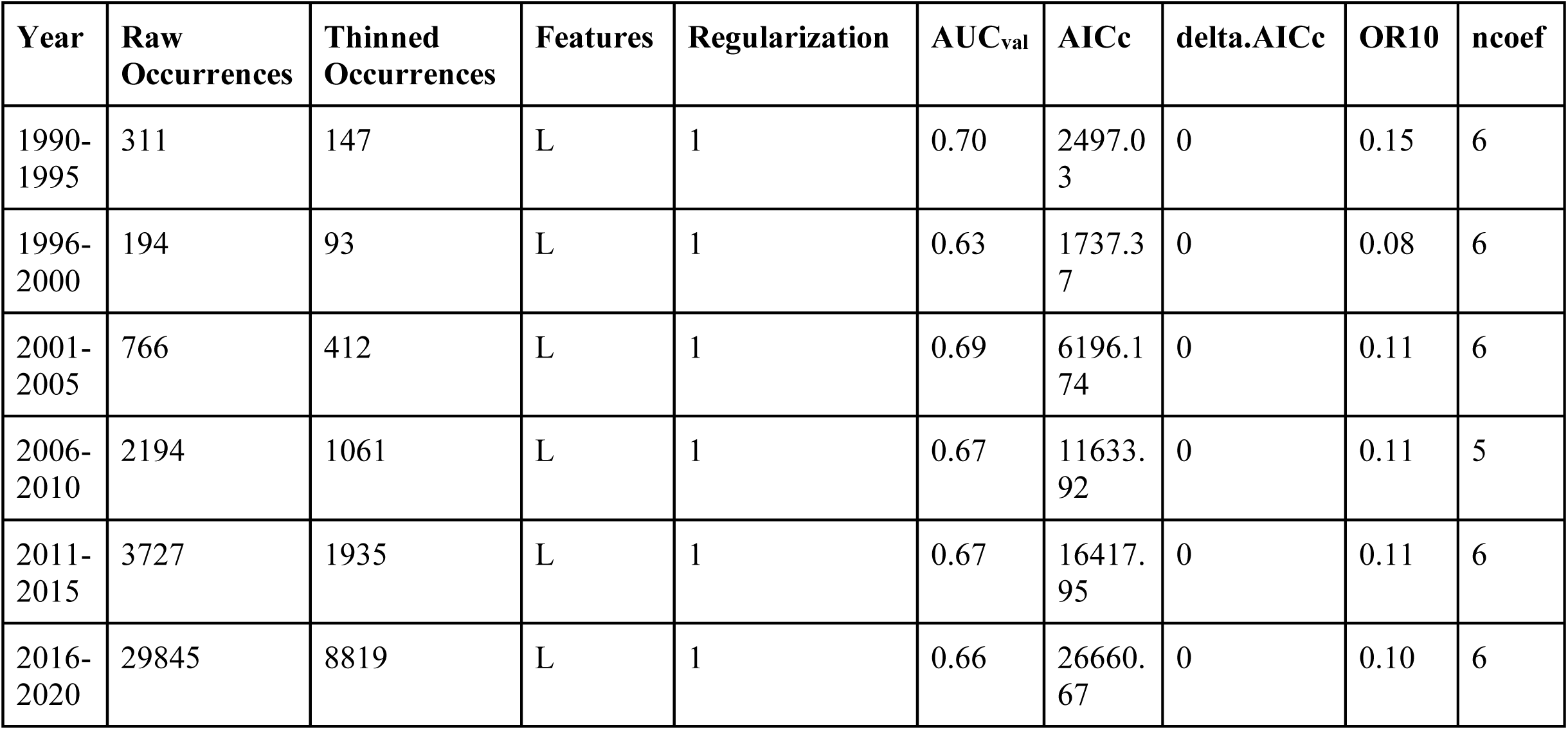
Sample size of occurrence localities before and after spatial thinning and maxent ENM settings for P. longipennis for the 5-yr intervals from 1990-2020. Settings shown are feature classes (features). Statistics shown are mean validation AUC (AUCval), mean omission rates for 10 percentile training values (OR10), Akaike Information Criterion (AICc), and delta Akaike Information criterion (delta.AICc), and number of non-zero coefficients (ncoef).

**Figure 3:**
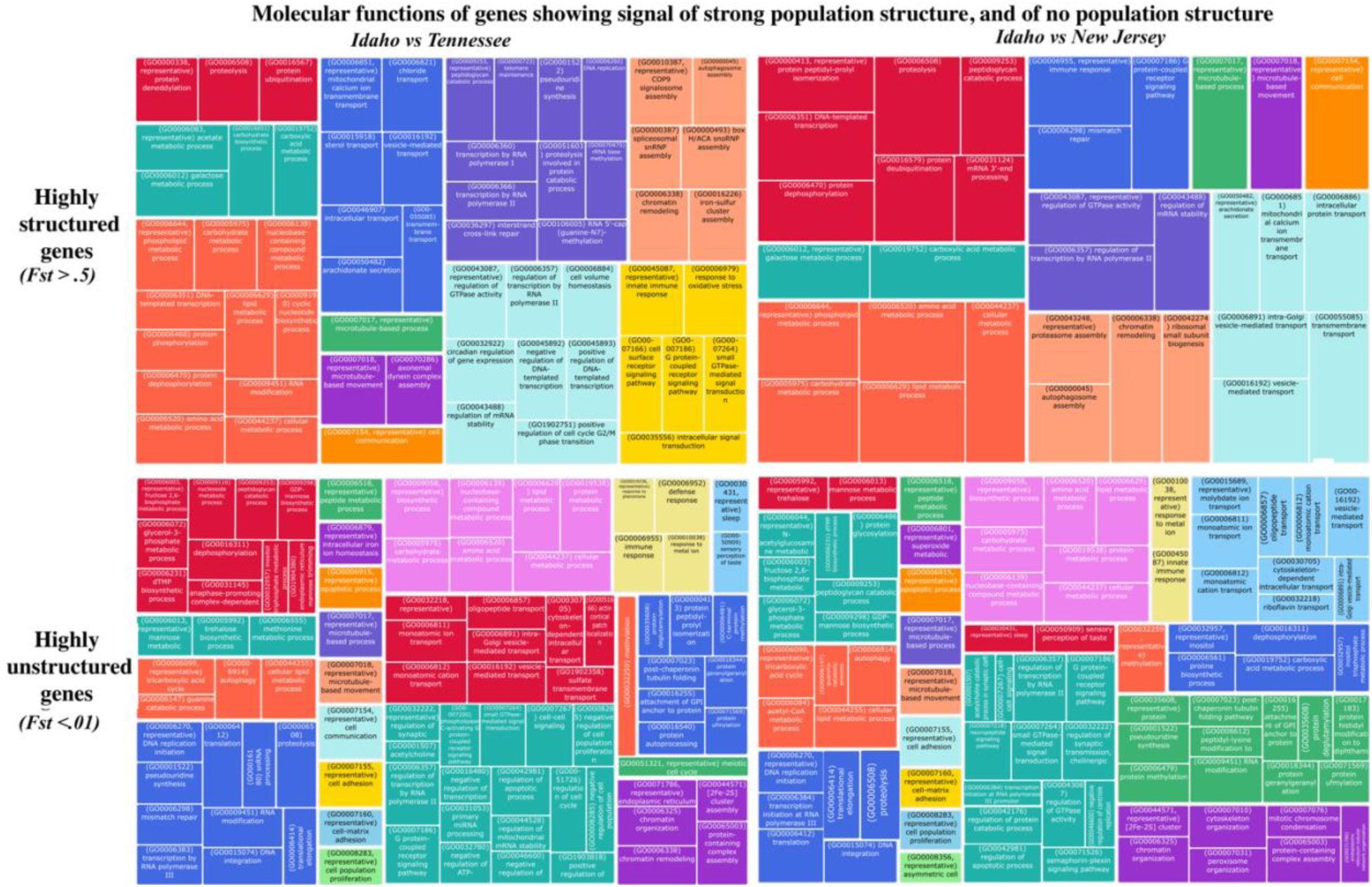
Molecular functions of genes from highly segregating (Fst >.5) and highly unstructured (Fst <.01) regions of the genome, calculated pairwise between the Idaho and New Jersey and Idaho and Tennessee populations with a sliding window of 10kb, with a step of 1kb.

In the pairwise comparison between Idaho and New Jersey, ∼29% of genes were highly unstructured, ∼ 9% of genes were greatly diverged. The unstructured genes consisted of 240 biological processes, 77 cellular components, and 271 molecular functions (Fig. 3). The structured genes consisted of 57 biological processes, 24 cellular components, and 82 molecular functions (Fig. 3).

The demographic model estimated a split between Idaho and the two East Coast populations ∼750ka with Tennessee and New Jersey splitting ∼50 Ka (Fig. 4A). Since their divergence, Tennessee is estimated to have been the recipient of frequent gene flow from New Jersey, and Idaho was a recipient of gene flow from the ancestral East coast population ∼400,000 years ago, and has been a recipient of gene flow from both populations more recently, in the past ∼50k years (Fig. 4A). The model identified one small burst of gene flow from Idaho to New Jersey (Fig. 4A). In this analysis, the simulated site frequency spectra (SFS) reasonably approximated the observed (Fig. 4B).

**Figure 4:**
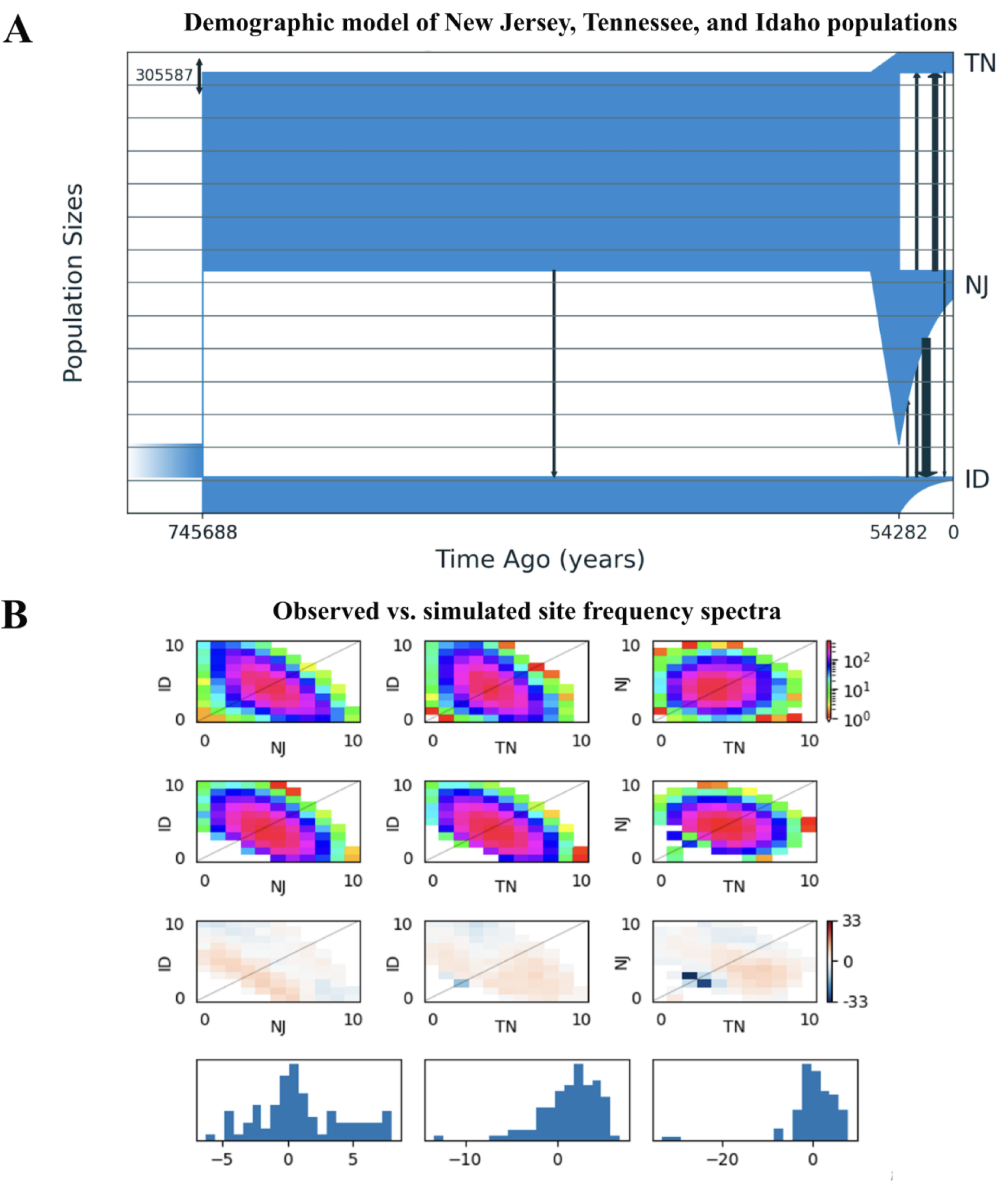
Historical Demography of P. longipennis. (A) Demographic model of the three P. longipennis populations as estimated by GADMA2 [35] demonstrating little contact between populations on either side of the Rocky Mountains following separation until bursts of gene flow occurred following the last glacial maxima. (B) Empirical and simulated pairwise site frequency spectra from the analysis.

### Morphological analysis

We were not able to successfully capture wing area, as different measurements of the same individual varied considerably due to a flaw in our semi-automated workflow that made it sensitive to scale bar variation (Supplementary Table 1). We found that the other wing measurements differed between the sexes when normalized by thorax length, but not when normalized by femur length (Table 2). When normalized by femur length the hindwing length, hindwing distal half, hindwing proximal half, hindwing aspect ratio, forewing length, forewing distal half, forewing proximal half and forewing aspect ratio all significantly differed between populations. The distribution of these measurements in the Idaho population skewed smaller than the other sites (Fig. 5), and Idaho was distal to the other populations across PC1 (loadings: hindwing length: 0.37; hindwind distal half: 0.37; forewing length: 0.37; hindwing depth: 0.36; forewing proximal half:.36), but not PC2 (loadings: hindwing depth: 0.82; forewing depth: 0.39; hindwing pterostigma: 0.36; forewing proximal half: 0.16, Fig. 5).

**Figure 5:**
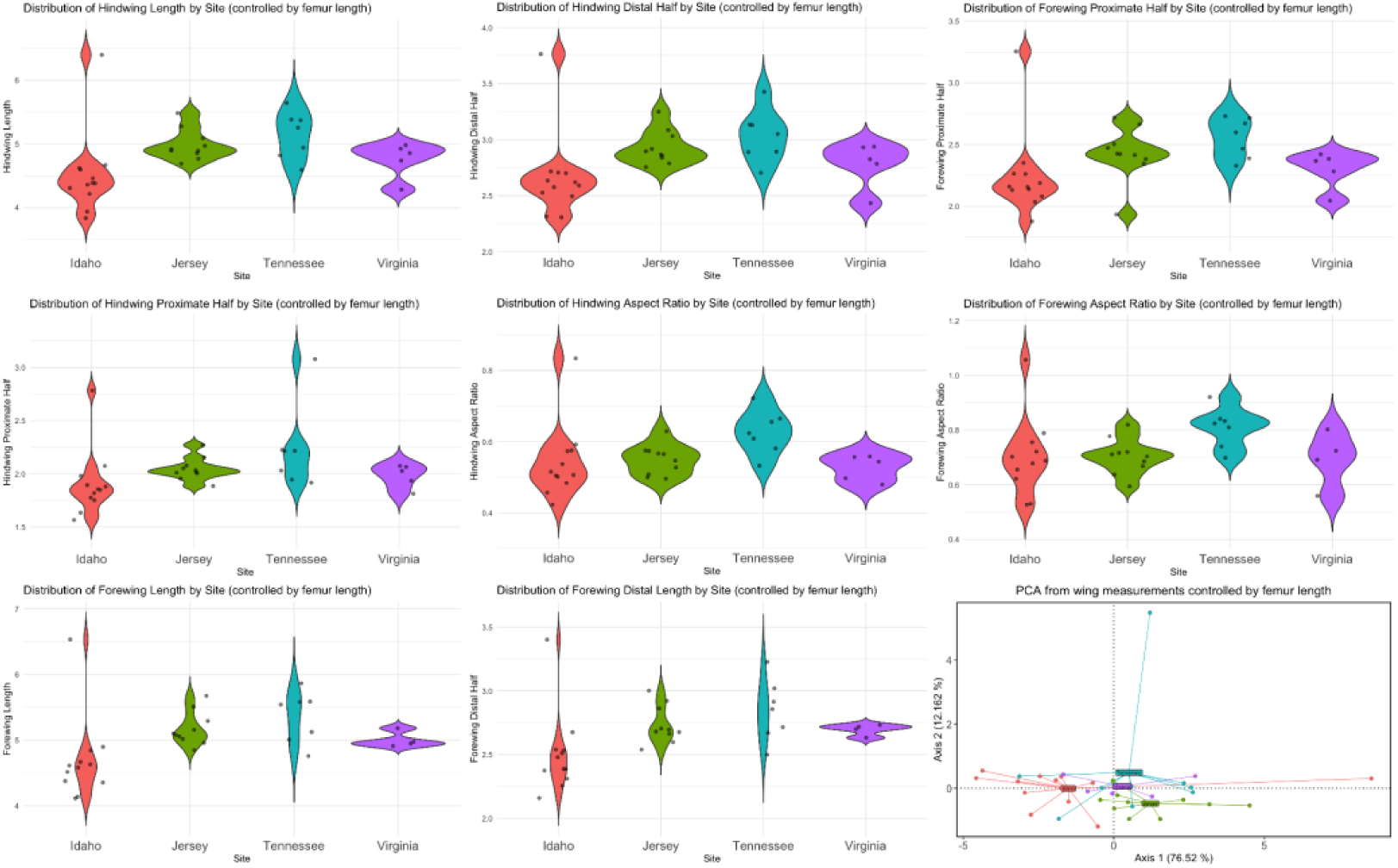
Wing measurements of P. longipennis. Violin plots of significantly differing wing measurements, when normalized by femur length, and PCA plot generated from wing measurements normalized by femur length.

### Ecological niche models

Optimal models exhibited robust results in 10-percentile omission rate but exhibited subpar results in mean validation AUC. Our mean validation AUC was low across time periods (0.63- 0.7), while 10-percentile omission rate was low (0.08-0.15) relative to its expected value of 0.1. Furthermore, models possessed a stark increase in AICc as the number of occurrences increased (AICc: 26660.67 for year 2016-2020).

Mean diurnal range (bio02) possessed the highest variable importance for the years 1990-1995, 2011- 2015, and 2016-2020 (range: ∼35-55%), maximum temperature of the warmest month (bio05) for the years 2006-2010 (∼35%), mean temperature of the warmest quarter (bio10) for the years 2001-2005 (∼50%), and precipitation of the driest month for the years 1996-2000 (∼34%). Mean diurnal range (bio02), temperature seasonality (bio04), maximum temperature of the warmest month (bio05), mean temperature of the warmest quarter (bio10), and precipitation of the driest month (bio14) possessed the second and third highest permutation importance throughout our models.

### Suitability Predictions

For the years 1990-1995, suitability was highest throughout the eastern United States, tapering off at the western half of Texas, with a pocket of suitability occurring in the Pacific Northwest (Fig. 6). For the years 1996-2000, suitability was highest along the southern and western coasts of the US (Fig. 6). For the years 2001-2005, suitability moves back to the entire eastern US, excepting Maine (Fig. 6). Throughout the years 2006-2020, suitability remains consistent, with the highest suitability along the southern and eastern coasts of the US, as well as the eastern US (Fig. 6).

**Fig. 6:**
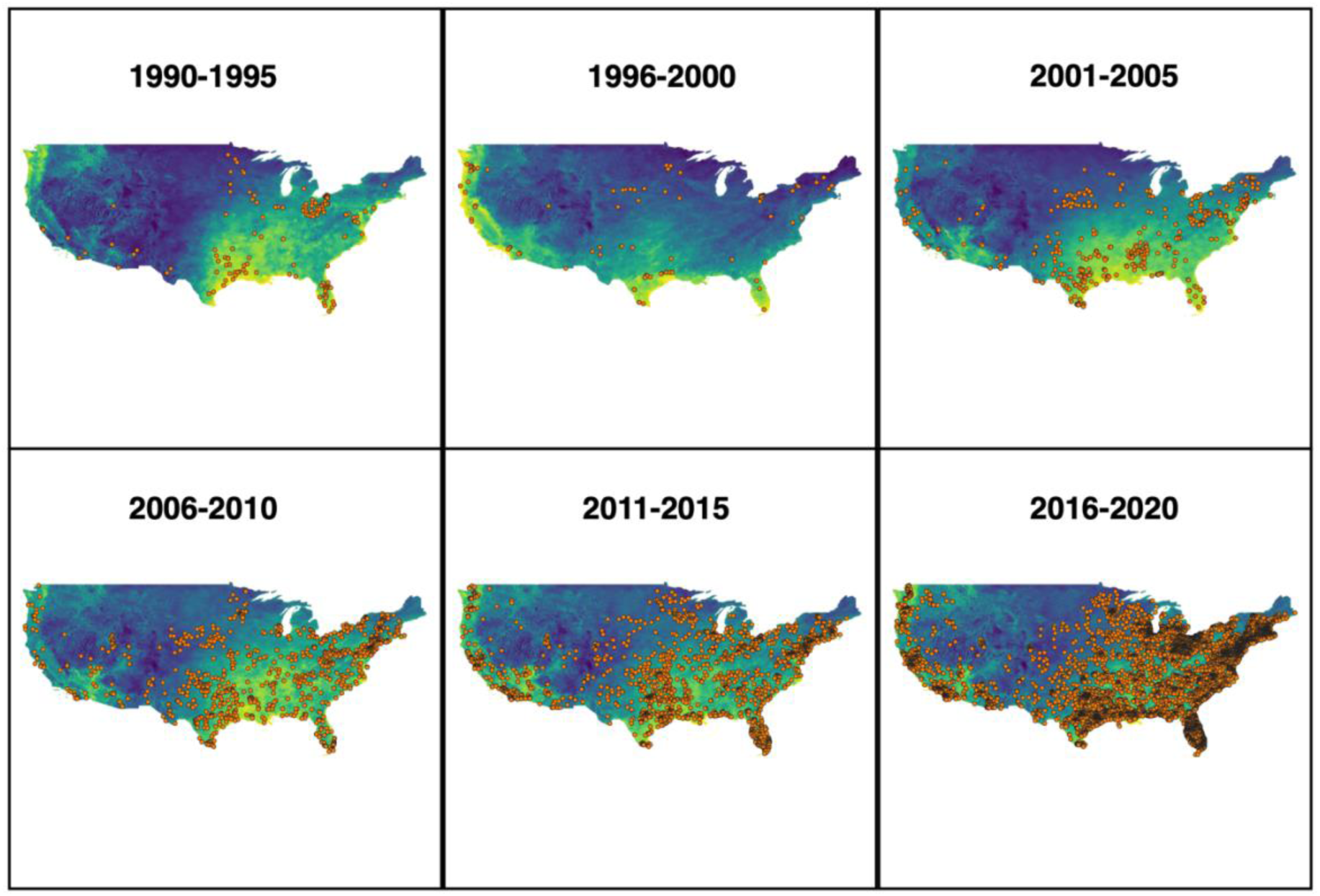
Maxent predictions for P. longipennis from the years 1990-2020, subdivided into 5 year time intervals. Predictions were derived using linear feature classes and 1x regularization. Predictions were transformed using the ‘cloglog’ function and converted to a range of 0-1 to approximate a probability of occurrence (Phillips et al., 2017). Brighter colors (yellow, green, blue) indicate areas of higher suitability (higher probability of occurrence), while darker colors (violet) indicate areas of lower suitability (lower probability of occurrence). Orange points are our thinned dataset for P. longipennis for each time slice.

## Discussion

### Dispersal capabilities of Pachydiplax

Our findings support the hypothesis that *P. longipennis* has a high dispersal capability, but it is not completely panmictic, as some migratory dragonflies are thought to be [65]. The two sampled East Coast populations are genetically quite similar (Fig. 2A-D, table 1), even more so in coding regions, and the demographic model shows some structure between Tennessee and New Jersey, but with a high degree of gene flow. Genetically, the population from Idaho does not appear to be a part of a migratory complex. It is genetically distinct from New Jersey and Tennessee (Fig. 2A-D, Table 1), and has primarily served as a sink population (Fig. 4A), all providing evidence that it is not dispersing towards eastern populations.

COI analyses also provided evidence of higher dispersal in eastern populations, which largely shared one COI haplotype, compared to western populations which were more disjointed (Fig. 2E).

Morphologically, the comparatively longer wings of the eastern populations may be related to selection for lower energy expenditure during gliding style flight [43], a possible adaptation for long distance travel. The molecular functions summarized from genes in highly structured regions between Idaho and Tennessee and Idaho and New Jersey provide evidence that Tennessee and New Jersey are equipped for longer-duration flight than *P. longipennis* from Idaho. Highly structured genes involved in the metabolism of carbohydrates and lipids, both of which are under heavy selection in migratory insects [66–68], and the selective pressures of high dispersal could be driven by positive selection associated with differing dispersal capabilities in *P. longipennis*, explaining the patterns observed in our F_ST_ scans (Fig. 3).

Much work remains to fully understand the dispersal patterns of *P. longipennis*. Broader sampling is needed to determine how *P. longipennis* dispersed to the Caribbean Islands and Bermuda, and the speed at which it should be expected to colonize newly available habitats. As a common pond dweller, it is possible that humans are transporting eggs and larvae with pond substrate, which could impact our estimates of gene flow and divergence times and facilitate dispersal to new habitats, but it cannot be ruled out that *P. longipennis* arrived at these islands through a natural dispersal process not directly involving human activity. Further genome wide resequencing data is needed to confirm the broad East/West population structure identified with COI, and to determine if any populations transverse the relatively inhospitable Rocky Mountains (Fig. 6).

### Molecular and morphological barriers between populations

Our molecular and morphological analyses revealed potential reproductive barriers between Idaho and the sampled eastern populations. One of the biological functions summarized from genes in genomic windows of high structure between Idaho and Tennessee was “circadian regulation of gene expression” (Fig. 3). Such genes have been implicated in temporal reproductive isolation in insects such as *Drosophila* [69], melon flies [70], Lepidoptera [71,72] and aphids [73], but no such work has been done in Odonata. Also of note are genes involved in “immune response” and “innate immune response” which segregate between Idaho and East Coast populations. In insects, immune response can have a strong influence on sexual reproduction and behavior [74,75].

Morphologically, it has been demonstrated that mate guarding, a behavior undertaken by *P. longipennis* [6] where a male guards the female until she deposits eggs (Fig. 2F), has a heavy effect upon front wings; indeed, forewings may be broader in species that utilize tandem mate guarding where males and females fly in copula to oviposit [76]. As female blue dashers on the East Coast have evolved with the longer-winged males (Fig. 5), it is unclear if the shorter-winged males from Idaho could carry out this behavior with East Coast females.

The implications of these findings on the species delimitation in the genus *Pachydiplax* should be further explored with more robust population and phenomic sampling.

### P. longipennis as an urban “extraordinaire”

#### Difficulties in modeling an abundant species

Our work confirms the hypothesis that *P. longipennis* is very well adapted to life in urban habitats. With the increased popularity of citizen science datasets starting approximately in 2010, the number of sightings of common species, like *P. longipennis*, increased dramatically. Indeed, as of December 2024, in its range, *P. longipennis* was the most widely observed aquatic insect on iNaturalist, the fifth most observed insect, and the 40th most observed species on iNaturalist. Although such increases in data provide the opportunity for highly robust niche models, a consequence is the overfitting of model performance due to the widespread abundance and density of *P. longipennis*. Predicted ENMs of *P. longipennis* revealed low AUC values across time slices. AUC values are a continuous metric, which provide discriminative ability in discerning between false positives and false negatives. Since *P. longipennis* is found throughout the continental United States, our models were unable to discern gradients of high and low suitability. This pattern is also observed in our AICc values. For the years 2016-2020, our thinned dataset consisted of 8,000+ occurrences. As a result, our AICc value was exceptionally high compared to the other models. This same observation can be seen on islands where occurrences are high for singular species [77]. Interestingly, our 10-percentile omission rate metrics were highly robust as most values were close to the expected value of 0.1. The 10-percentile omission rate metric excludes the lowest 10 percent of raw occurrence values, thus creating a threshold within suitability. Our low omission and AUC values suggest minimal outliers in our dataset, suggesting that our predictions are not overly generalist.

#### Adaptations for success in human altered environments

Our models estimate that extremes in temperature, precipitation, and temperature fluctuation are the main drivers of the response of *P. longipennis* to their distribution. Due to their widespread abundance throughout the contiguous United States, it seems this species has an exceptionally high thermal tolerance and desiccation threshold, which elucidates why they are so widespread in urban environments. In reared experiments, larval development of *P. longipennis* was unaffected by increases in temperature or resource level [78]. Furthermore, mesocosm experiments [79] revealed that larval *P. longipennis* are more likely to engage in intraguild predation with increases in temperature, which may allow them to diversify prey types in warmer habitats. Urban environments have significantly higher ground temperatures than rural areas, a consequence of the higher demand of power plants providing energy for in-home cooling units and impervious surfaces creating urban heat islands [80]. We propose that *P. longipennis* is distributed widely in urban environments due to their optimal habitats being warmer, a condition that will only increase as the urban heat sink phenomenon continues to worsen.

Our suitability predictions suggest that the distribution of *P. longipennis* is widening as time progresses and urbanization increases throughout the United States. However, one caveat of our distribution sampling is the impact of the increase of documented *P. longipennis* sightings as the result of increasingly popular community science programs. Although our models may be slightly skewed towards increases in distribution of *P. longipennis*, our genomic analyses support the biological realism of the models. One molecular function previously identified in expanding gene families in the genome assembly of *P. longipennis* was the “Peptidoglycan Catabolic Process” [4]. Peptidoglycan is an integral compound in most bacterial cell walls [81], and genes associated with peptidoglycan catabolism are thought to be involved in both immune and desiccation response in insects [82]. It is intriguing that between Idaho and the East Coast populations of *P. longipennis,* examples of highly structured and highly unstructured genes can *both* be tied to this molecular process (Fig. 3). Our population genomics analysis provides further evidence that these gene families play a key role in the dispersal success of *P. longipennis* in polluted habitats and are a driver of the expanding niche of *P. longipennis*. In addition to enabling *P. longipennis* to survive amidst the higher desiccation risk found in warmer habitats, the lentic water bodies they inhabit are expected to carry a high bacterial load. It is possible that certain peptidoglycan variants will be encountered by all populations of *P. longipennis*, explaining unstructured genes related to these functions. Each population of *P. longipennis* could also encounter unique variants of peptidoglycan, explaining the high segregation in some of these genes. Such a dynamic could be extended to an identical phenomenon observed involving genes which are associated with “immune response” and “innate immune response.” Sampling of *P. longipennis* alongside bacterial communities will be an important step in testing this hypothesis.

Furthermore, *P. longipennis* likely faces high levels of exposure to free radicals, similar to other animals in urban environments [83]. Expanded gene families involved in the processing of free radicals have been hypothesized as an important mechanism enabling P. longipennis to successfully colonize urban environments [33]. Our finding that genes involved in this process are highly unstructured between populations (Fig. 3) provides further evidence that these gene families are key to the success of *P. longipennis*, in addition to its ability to respond to desiccation and a high pathogen load. Future research with these genes will likely include transcriptome sequencing to understand patterns of regulation in association with oxidative stress.

Recent niche models of odonates have suggested high-dispersing species will be the ecological “winners” in a changing global climate [84–86]. It seems that *P. longipennis* is definitively a winner that will flourish as humans expand. We do not expect the genes involved in the success of *P. longipennis* in urban environments to be restricted to this species. Further work needs to be done in other such species to determine if there is convergence in species that inhabit these environments.

## Conclusion

Our work presents compelling evidence that *P. longipennis* is a successful habitat generalist. Studies of lentic habitats in California between the early Twentieth and early Twenty-first Centuries found a shift in odonate assemblages away from specialists and towards generalists [7]. Among the species showing the greatest expansion was *P. longipennis*. This study confirms that *P. longipennis* is well equipped to deal with the stress of urban habitats. East Coast populations have a relatively higher dispersal capability (Fig. 2), which could allow them to more rapidly colonize newly suitable habitat. While the specific dynamics between bacterial loads, oxidative stress, and the establishment *P. longipennis* need further study we posit that the dominance of the highly observable *P. longipennis* at a water body can be considered evidence of contamination and biodiversity loss at a given habitat.

## Acknowledgements

We would like to thank Dr. Michael L. May for his foundational work on *P. longipennis*, which made this study possible. We would also like to thank Dr. Anton Suvorov for his support. This work was supported by start-up funding to MKK from Baruch College, CUNY, and the Boise Youth Climate Action Fund.

## Data availability

The “Gadma” parameters file, rMarkdown script used for statistical analysis, images, and phyllip file use for COI analyses are available on figshare (10.6084/m9.figshare.28020914). The raw sequencing reads have been uploaded to the sequence read archive (bioproject PRJNA1199395).

## Conflicts of interest

The authors declare no conflicts of interest.

## Notes

https://figshare.com/articles/dataset/Supplementary_materials/28020914

### Competing Interest Statement

The authors have declared no competing interest.

### Summary of Updates

The manuscript has been updated to reflect BioProject assignment by NCBI.

