## Supplementary Materials for "The Blueprint for Survival: The Blue Dasher Dragonfly as a Model for Urban Adaptation"

RU–N:WL:OdoD: 2013 00821
Ware Lab Specify Database
Rutgers University,Newark,NJ

AMERICAN MUSEUM of NATURAL HISTORY

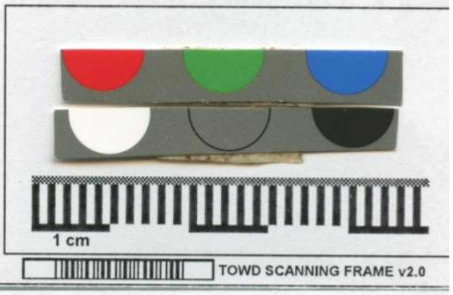


Pachydiplax longipennis ♂

NJ: Passaic Co, Garret Mtn
Res, Barbour Pond
40.8991°N, 74.1831° W
12-VII-2012 Coll: Aim High

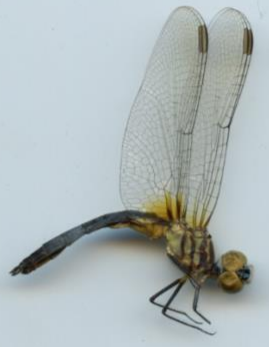

*Supplementary figure 1: Scanned specimen using TOWD template*

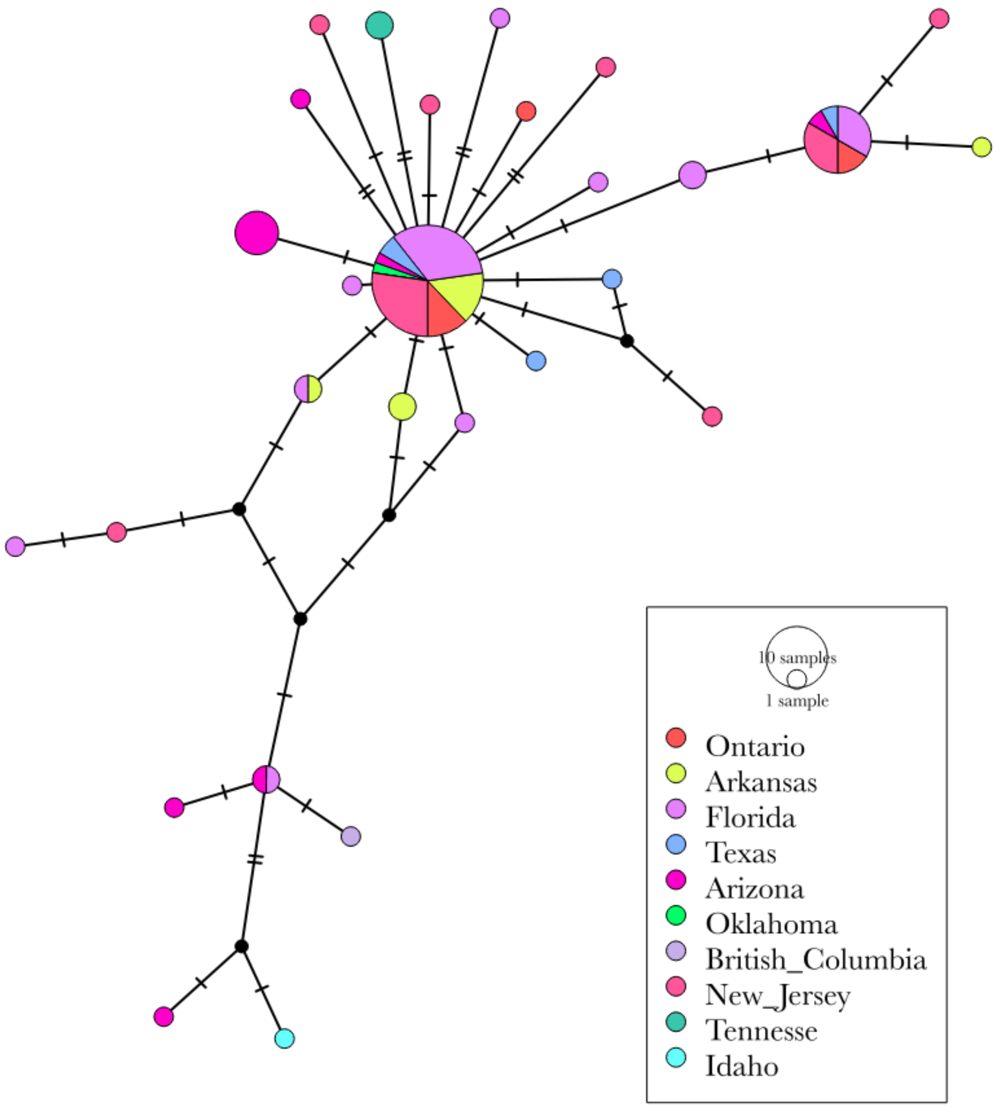


*Supplementary Figure 2: COI haplotype network of P. longipennis, including sequences from Tennessee*

| Supplementary Table 1: Wing Area Measurements | | | | |
|---|---|---|---|---|
| image_name | Individual | Population | hindwing_area_cm2 | forewing_area_cm2 |
| 0AC1C123-FA41-4C15-8E0F-B2551A24B2C7_1_105_c.jpeg | 1 | Idaho | 1.754010642 | 1.215001684 |
| 5061AF63-4C1C-42CB-8AEB-04EC3785B20F_1_105_c.jpeg | 1 | Idaho | 1.966922516 | 1.368680454 |
| 8BC3CEF6-CA23-431F-8BB9-6E61636EEA01_1_105_c.jpeg | 2 | Idaho | 1.953961845 | 1.244922116 |
| 7CC6F43E-0F82-488B-93A5-626AF1BFB980_1_105_c.jpeg | 2 | Idaho | 1.567989906 | 1.284168587 |
| B97D7ED4-6E26-450E-8F0D-542A481EA108_1_105_c.jpeg | 3 | Idaho | 2.14006918 | 1.678566184 |
| 6B388BE7-A825-4775-8756-B38E432C0156_1_105_c.jpeg | 3 | Idaho | 1.556571816 | 1.139464192 |
| 7FE42C93-C07A-4EA9-8780-0741FF8996DE_1_105_c.jpeg | 3 | Idaho | 2.133115092 | 1.730003736 |
| BAAD116A-6FF0-4D79-8232-559A77EEA9E5_1_105_c.jpeg | 4 | Idaho | 2.116299741 | 1.483539078 |
| BB779614-0E51-4E2B-AA67-A43A93ADC69E_1_105_c.jpeg | 4 | Idaho | 2.399309145 | 2.001636824 |
| D8FF208C-0365-4618-B578-3572371DA4AB_1_105_c.jpeg | 4 | Idaho | 1.667347863 | 1.479162816 |
| 3B2CB93D-4E11-4A03-9C16-D9FF88B407A6_1_105_c.jpeg | 4 | Idaho | 2.180012313 | 1.804404222 |
| BD19F957-3660-43E8-98CA-74E772288A60_1_105_c.jpeg | 5 | Idaho | 1.746145446 | 1.234471241 |

| Supplementary Table 1: Wing Area Measurements | | | | |
|---|---|---|---|---|
| D4426030-E40A-44EF-8A8F-A741754177C2_1_105_c.jpeg | 5 | Idaho | 2.358871531 | 1.841295545 |
| 1EA076EE-4586-4EF7-B5EF-6DAF1350577D_1_105_c.jpeg | 5 | Idaho | 1.674049811 | 1.124865905 |
| 3D32BD18-C953-4617-B9DC-818A63B5E5F9_1_105_c.jpeg | 5 | Idaho | 2.42782722 | 1.807239168 |
| 19A38A95-507E-475F-BCBA-66E82C6764FB_1_105_c.jpeg | 7 | Idaho | 1.711127201 | 1.366134228 |
| 9829F61C-2ABB-4353-8B90-A80101DD4AC4_1_105_c.jpeg | 7 | Idaho | 2.320771169 | 2.106510907 |
